# An indispensable role for dynamin-related protein 1 (DRP1) in adipogenesis

**DOI:** 10.1101/2020.04.13.039149

**Authors:** Raja Gopal Reddy Mooli, Dhanunjay Mukhi, Zhonghe Chen, Nia Buckner, Sadeesh K. Ramakrishnan

## Abstract

Emerging evidence indicates that proper mitochondrial dynamics is critical for adipocyte differentiation and functional thermogenic capacity. We found that mitochondrial fission protein dynamin-related protein 1 (DRP1) is highly expressed in brown adipose tissue compared to white adipose tissue and their levels increase during brown adipocyte differentiation. Our results reveal that the inhibition of DRP1 using Mdivi-1 mitigates adipocyte differentiation and differentiation-associated mitochondrial biogenesis. We found that DRP1 is essential for the induction of the early phase of adipogenic transcriptional program. Intriguingly, inhibition of DRP1 is dispensable following the induction of adipogenesis and adipogenesis-associated mitochondrial biogenesis. Together, we demonstrate that DRP1 in the preadipocytes plays an essential role in brown and beige adipogenesis.

## Introduction

The adipose tissue acts as a central organ in regulating metabolic homeostasis and is broadly categorized into two types; white adipose tissue (WAT) and classical brown adipose tissue (BAT) (Choe, Huh, Hwang, Kim, & Kim, 2016; Kahn, Wang, & Lee, 2019). Recent studies have uncovered an inducible form of adipocytes in white fat depots namely beige (or brite) adipocytes (Ikeda, Maretich, & Kajimura, 2018; Kajimura, Spiegelman, & Seale, 2015; Wu et al., 2012). Beige adipocytes are induced by cold exposure, β3-selective adrenergic agonists and by dietary intervention such as caloric restriction (Barbatelli et al., 2010). White adipocytes store energy storage as triglycerides while the beige and brown adipocytes expend energy via non-shivering thermogenesis (Giralt & Villarroya, 2013; Sidossis & Kajimura, 2015). The metabolic distinction between the adipocytes is partly driven by their mitochondrial characteristics. For example, white adipocytes have fewer mitochondria limiting their oxidative capacity, whereas brown and beige adipocytes possess abundant mitochondria expressing uncoupling protein 1 (UCP1) involved in non-shivering thermogenesis (Sepa-Kishi & Ceddia, 2018; Vosselman, van Marken Lichtenbelt, & Schrauwen, 2013). Thus, the mechanisms associated with mitochondrial remodeling have a key role in regulating adipocyte function.

Mitochondrial remodeling is tightly regulated by the mitochondrial dynamics which includes fission and fusion (Scott & Youle, 2010; Tilokani, Nagashima, Paupe, & Prudent, 2018). Fission is mediated by the cytosolic protein, dynamin-related protein 1 (DRP1), which is recruited to the mitochondrial surface with the help of accessory proteins such as mitochondrial fission factor (MFF) and mitochondrial fission protein 1 (FIS1) (Bleazard et al., 1999). Fission is followed by fusion, which is coordinated by the inner and outer mitochondrial proteins such as mitofusin (MFN)-1 and -2 and optic atrophy 1 (OPA1) (Westermann, 2002; Zhang & Chan, 2007). Studies have shown that the dysregulation of mitochondrial dynamics has a profound effect on cellular energetics which in turn could affect their function (Eisner, Picard, & Hajnoczky, 2018; Mishra & Chan, 2016). For instance, impairment in mitochondrial fragmentation decreases the reprogramming and differentiation efficiency of the progenitor cells (Forni, Peloggia, Trudeau, Shirihai, & Kowaltowski, 2016). Similarly, inhibition of DRP1-mediated mitochondrial fragmentation mitigates the self-renewal potency or stemness of the progenitor cells (Khacho et al., 2016). Thus, mitochondrial dynamics plays a key role in governing cellular property and function.

Although studies have highlighted the importance of adipose tissue mitochondrial dynamics from a metabolic perspective, very little is known about its contribution to adipogenesis. We found that DRP1 is highly expressed in the BAT and their levels increase during adipocyte differentiation. Here, we report that inhibition of DRP1 during the early phase of differentiation attenuates adipogenesis and adipogenesis-induced mitochondrial biogenesis. Thus, we demonstrate an indispensable role of DRP1 in regulating adipocyte differentiation.

## Results

### DRP1 is highly expressed in brown and beige adipocytes

DRP1 plays an essential role in regulating mitochondrial homeostasis by inducing mitochondrial fission (Bleazard et al., 1999). Since brown adipocytes possess an abundance of mitochondria, we explored the expression of DRP1 by Western blot analysis. Our data shows that the brown adipose tissue (BAT) express high levels of DRP1 when compared to the inguinal white adipose tissue (iWAT) or epididymal white adipose tissue (eWAT) (Figure 1A). Since beige adipocytes have mitochondrial characteristics similar to brown adipocytes, we further determined whether beigeing increases DRP1 expression. To this end, C57BL6 mice were injected with β3-AR agonist (CL316, 243; CL) for 7 consecutive days, which induced beige adipocytes as revealed by an increase in the expression of uncoupled protein 1 (UCP1) in the iWAT. We found that beigeing was associated with an increase in DRP1 in the iWAT (Figure 1B). We then assessed the expression of DRP1 in the transformed mouse stromal vascular fraction (SVF) cells that are differentiated into brown adipocytes. The expression of DRP1 increased as early as 24 hours of induction and found a time-dependent increase in their expression during adipocyte differentiation (Figure 1C). qPCR analysis showed a parallel increase in *Drp1* mRNA levels during adipocyte differentiation (Figure 1D). Similar to the transformed SVF cells, differentiation of adipocyte precursor SVF cells increased DRP1 expression (Figure 1E). When we further compared the expression of DRP1 between preadipocytes differentiated to brown or white adipocytes, DRP1 levels were higher in the differentiated brown adipocytes when compared to the white adipocytes (Figure 1F). Together, our data indicate that DRP1 is highly expressed in the brown and beige adipocytes.

**Figure 1.**
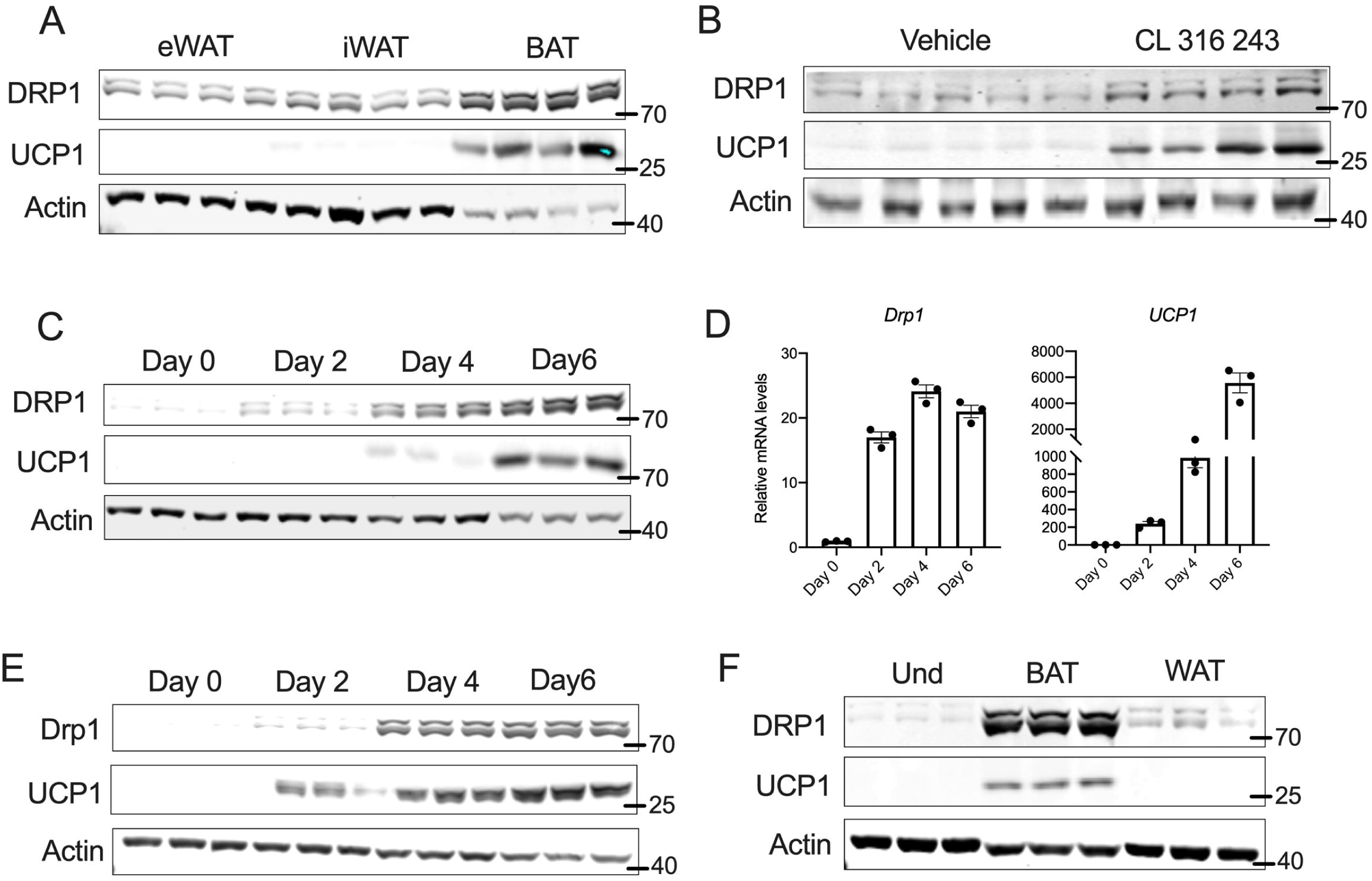
DRP1 is highly expressed in brown and beige adipocytes. (A) Immunoblots of DRP1 and UCP1 in various fat depots of 6-week old C57BL6 mice. (B) Immunoblots of DRP1 and UCP1 in the subcutaneous fat of C57BL6 mice injected with β3-adrenergic agonist CL316, 243 intraperitoneally at a dose of 1mg/kg bodyweight for 7-days. (C) Immunoblots for DRP1 and UCP1 in transformed mouse stromal vascular fraction (SVF) cells differentiated into brown adipocytes. (D) mRNA levels of DRP1 and UCP1 in transformed mouse SVF cells differentiated into brown adipocytes. β-actin was used to normalize the gene expression. (E) Immunoblots for DRP1 and UCP1 in primary mouse SVF cells from subcutaneous fat of C57BL6 that are differentiated into brown adipocytes. (F) Immunoblots for DRP1 and UCP1 in transformed mouse SVF cells differentiated into brown or white adipocytes. Und-Undifferentiated

### DRP1 is essential for the thermogenic program of the brown adipocytes

The activity of DRP1 is determined by its GTPase domain, which promotes DRP1 self-assembly into oligomers (Smirnova, Griparic, Shurland, & van der Bliek, 2001). To investigate the role of DRP1 in adipogenesis, we differentiated the SVF cells into brown adipocytes in the presence or absence of Mdivi-1, an inhibitor of DRP1 GTPase activity (Bleazard et al., 1999). Inhibition of DRP1 from day 0 of differentiation abolished the induction of the UCP1 (Figure 2A). PRDM16, a large zinc finger-containing transcription factor, is the master regulator of thermogenic gene expression (Harms et al., 2014). Inhibition of DRP1 decreased the expression of PRDM16 (Figure 2A). Moreover, the mRNA levels of other brown fat-specific genes such as *Ucp1, Prdm16, Cidea*, and *Dio2* in the adipocytes differentiated in the presence of Mdivi-1 (Figure 2B).

**Figure 2.**
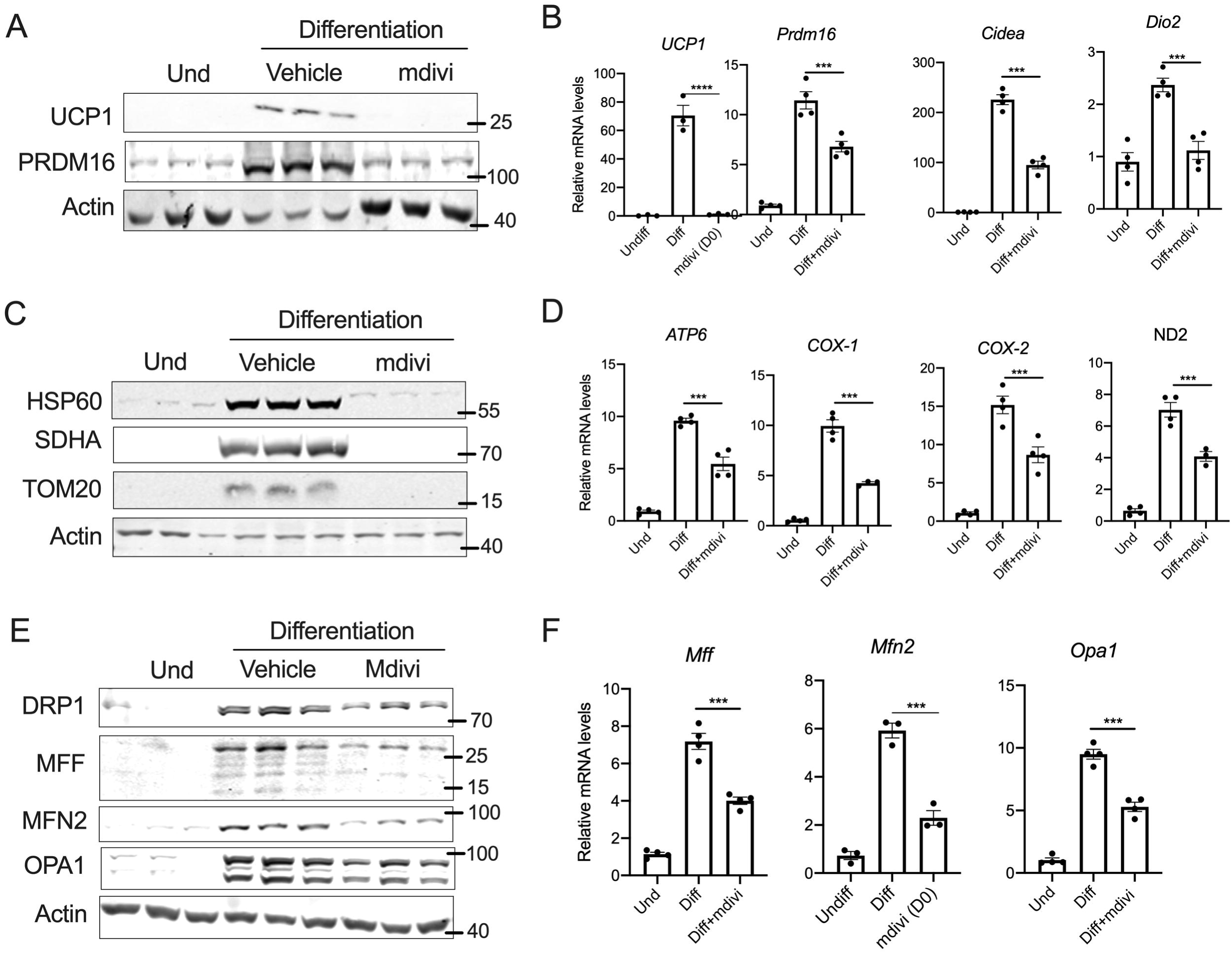
DRP1 is essential for the thermogenic program of the brown adipocytes. (A) Immunoblots and (B) qPCR analysis of thermogenic genes in the SVF cells differentiated to brown adipocytes in the presence or absence of 10μM Mdivi-1 added from day 0 of differentiation. (C) Immunoblots and (D) qPCR analysis for the mitochondrial genes in the SVF cells differentiated to brown adipocytes in the presence or absence of 10μM Mdivi-1 added from day 0 of differentiation. (E) Immunoblots and (F) qPCR analysis of mitochondrial dynamics-related genes in the SVF cells differentiated to brown adipocytes in the presence or absence of 10μM Mdivi-1 added from day 0 of differentiation. β-actin was used to normalize the gene expression. The results are presented as SE ± SEM (***p < 0.001). Und-Undifferentiated

Brown adipocyte differentiation is strongly associated with mitochondrial biogenesis (Uldry et al., 2006). Since DRP1-mediated fission is part of the mitochondrial biogenesis machinery and the thermogenic program is strongly associated with mitochondrial biogenesis (Cannon & Nedergaard, 2008), we assessed whether inhibition of DRP1 affects adipogenesis-mediated mitochondrial biogenesis. Western blot analysis showed that the expression of mitochondrial proteins such as HSP60, SDHA and TOM20 were decreased in the adipocytes differentiated in the presence of Mdivi-1(Figure 2C). Besides, Mdivi-1 treatment decreased the mRNA levels of the numerous mitochondrial genes such as *Atp6, Cox-1, Cox-2*, and *Nd2* (Figure 2D). When we further assessed for the mitochondrial dynamics-related proteins, the expression of MFF, OPA1, and MFN2, were decreased both at the mRNA and protein levels in the Mdivi-1 treated adipocytes (Figure 2E and 2F). Together, the data demonstrate that inhibition of DRP1 mitigates the thermogenic mitochondrial remodeling of the brown adipocytes.

### DRP1 inhibition ablated the adipogenic program

The morphological assessment of differentiated adipocytes showed a dramatic decrease in lipid droplets when DRP1 is inhibited (Figure 3A). Therefore, we investigated whether DRP1 inhibition attenuated adipocyte differentiation, which is regulated by a complex network of transcriptional factors (Farmer, 2006). During the early phase of adipocyte differentiation, the activation of CCAAT/enhancer-binding proteins β and δ (C/EBPβ and C/EBPδ) is followed by the induction of C/EBPα and peroxisome proliferator-activated receptor γ (PPARγ) (Guo, Li, & Tang, 2015). The late phase of adipogenesis involves C/EBPα and PPARγ-mediated expression of PGC-1α and other genes involved in adipocyte differentiation (Guo et al., 2015; Rosen & MacDougald, 2006; Rosen, Walkey, Puigserver, & Spiegelman, 2000). Inhibition of DRP1 from day 0 of differentiation decreased the expression of key transcription factors involved in both the early and late phases of adipocyte differentiation (Figure 3B). Consistently, the mRNA levels of various adipogenesis-specific genes showed a similar decrease in their expression upon DRP1 inhibition (Figure 3C). Collectively, the data shows that DRP1 plays an essential role in adipocyte differentiation.

**Figure 3.**
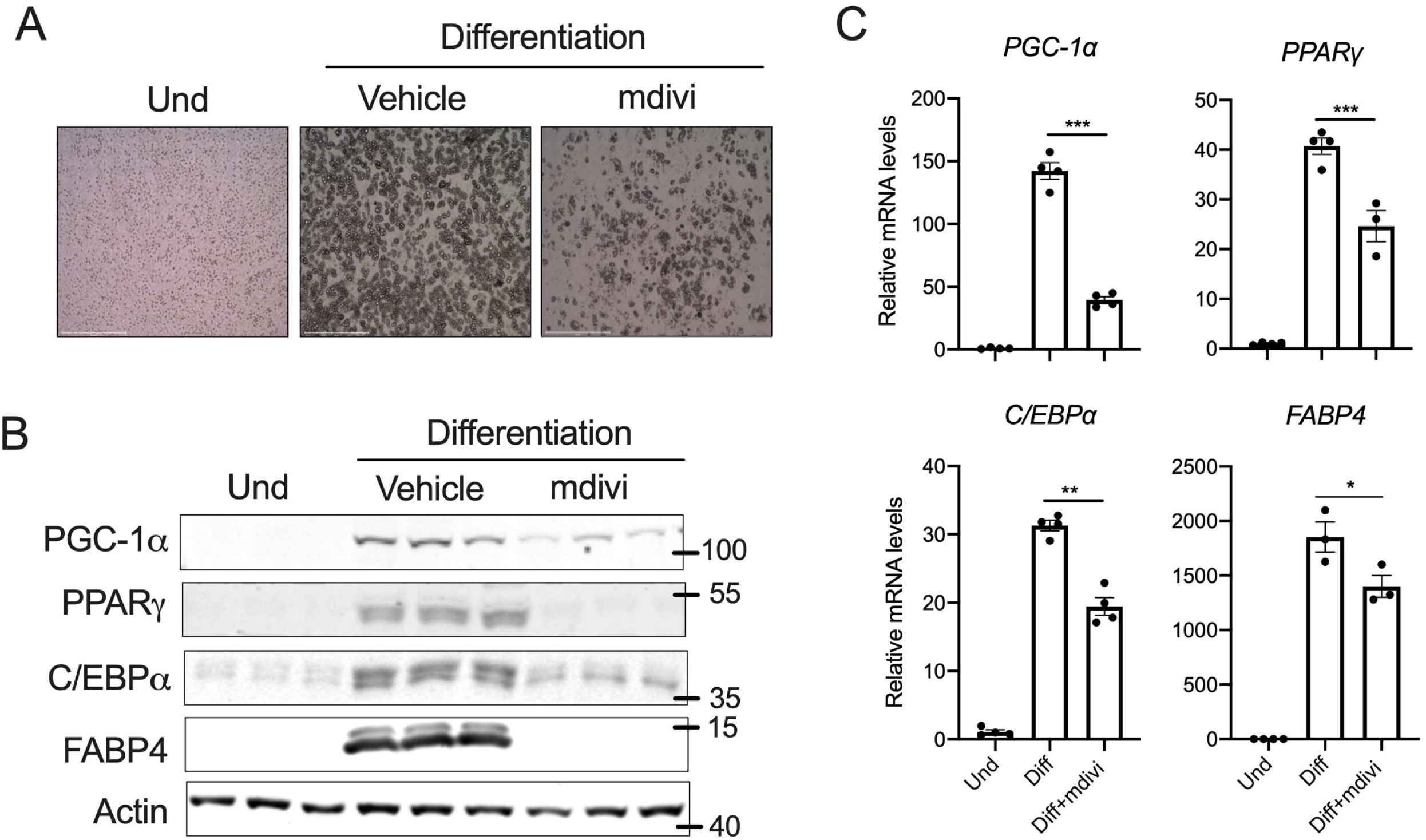
DRP1 inhibition attenuates the adipogenic program. (A) Representative images of the SVF cells differentiated to brown adipocytes in the presence or absence of 10μM Mdivi-1 added from day 0 of differentiation. (B) Immunoblots and (C) qPCR analysis of key transcription factors involved in adipogenesis in differentiated brown adipocytes in the presence or absence of 10μM Mdivi-1 added from day 0 of differentiation. β-actin was used to normalize the gene expression. The results are presented as SE ± SEM (*p < 0.05, **p < 0.01***, p < 0.001). Und-Undifferentiated

### DRP1 is indispensable for the induction of adipocyte differentiation

To further delineate the requirement of DRP1 for adipocyte differentiation, we inhibited DRP1 during the induction period alone by treating the SVF cells with Mdivi-1 on day 2 (D2) and then washed the wells with media and continued the (maintenance media) differentiation without Mdivi-1. Restoring DRP1 activity by washing out Mdivi-1 did not rescue the differentiation potential of the SVF cells as revealed by the morphological assessment for the lipid droplets (Figure 4A). Moreover, the expression of adipogenesis-related proteins was not restored in Mdivi-1 washout cells (Figure 4B) indicating that inhibiting DRP1 during the induction period is sufficient to suppress adipocyte differentiation. We then sought to determine the mechanism by which inhibition of DRP1 during the induction period attenuates adipogenesis. The expression of the adipogenesis-related genes (PGC-1α, PPARγ and C/EBPα) and mitochondria-related genes (Cox-1& 2, ND2, ATP6) were induced as early as 24-hours following differentiation (Figure 4C and 4D). We found that the inhibition of DRP1 significantly attenuated the early induction of adipogenesis- and mitochondria-related genes both at the mRNA and protein level (Figure 4E and 4F) suggesting that DRP1 plays a critical role in the transcriptional programming of adipocyte differentiation. Since a nuclear expression of DRP1 has been demonstrated (Chiang et al., 2009), we questioned whether the adipogenesis induces nuclear translocation of DRP1. However, no detectable levels of DRP1 were observed in the nucleus of SVF cells with or without differentiation (Figure 4G), suggesting that cytosolic DRP1 regulates the transcriptional programming of adipocyte differentiation.

**Figure 4.**
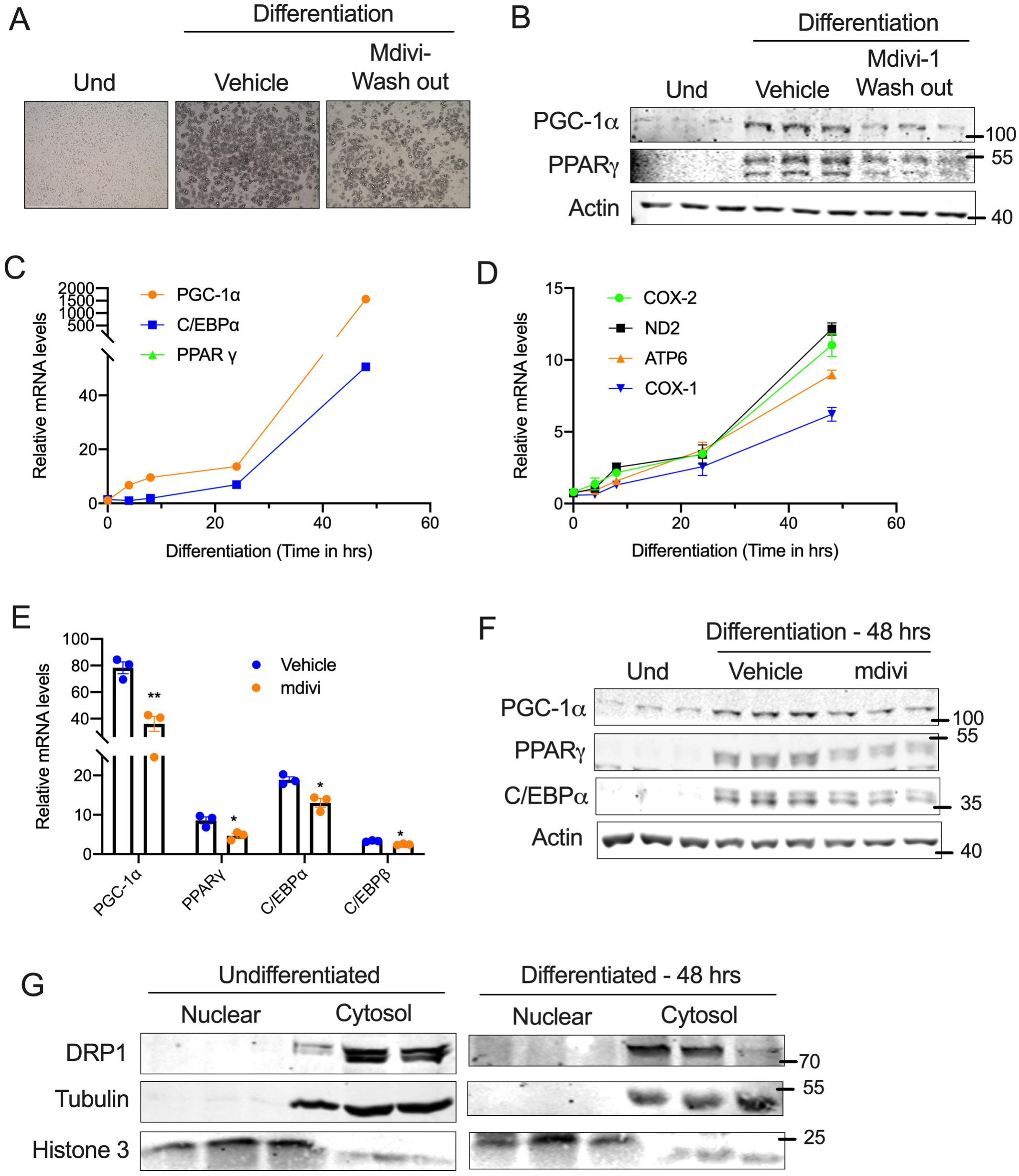
DRP1 is indispensable for the induction of adipocyte differentiation. (A) Representative images for the SVF cells differentiated to brown adipocytes in the absence or presence of 10μM Mdivi-1 during the induction time (48hrs) and then the wells were washed and continued on maintenance media without Mdivi-1 for the next 6 days. (B) Immunoblots for the adipogenesis-related proteins. qPCR analysis assessing the time-dependent change in the expression of genes involved in (C) adipogenesis and (D) mitochondrial markers during the early phase of adipocyte differentiation. (E) qPCR and (F) Western blot analysis for genes involved in adipogenesis in the SVF cells assessed at 48hrs following induction of differentiation. (G) Immunoblots of DRP1 in nuclear and cytosolic fractions of SVF cells assessed at 48hrs following induction of differentiation. β-actin was used to normalize the gene expression. Tubulin and histone 3 were used as cytosolic and nuclear loading control, respectively. The results are presented as means SE ± SEM (*p < 0.05, **p < 0.01, ***p < 0.001). Und-Undifferentiated

### DRP1 is dispensable during the late phase of adipocyte differentiation

Thus far, we demonstrate that inhibition of DRP1 during the course of differentiation suppressed adipocyte differentiation. The acquisition of mature adipocyte phenotype from precursor cells involves sequential steps, which is reflected by the appearance of early, intermediate and late mRNA/protein markers of adipocyte differentiation (Kelly & Scarpulla, 2004). To further elucidate the role of DRP1 in adipogenesis, we took an alternative approach where we inhibited DRP1 during the maintenance period alone. To this end, SVF cells were differentiated with a brown adipocyte cocktail for two days. On day 2, the cells were switched to the maintenance media with or without Mdivi-1 and continued with the respective treatments throughout the differentiation period. Interestingly, inhibition of DRP1 during the maintenance period (D2) did not affect adipogenesis as revealed by visual examination of lipid droplets (Figure 5A). Moreover, inhibition of DRP1 during the maintenance period did not affect the mRNA and protein levels of adipogenesis or mitochondria-related genes (Figure 5B-E). Collectively, the data emphasize that DRP1 is dispensable during the maintenance period of adipocyte differentiation.

**Figure 5.**
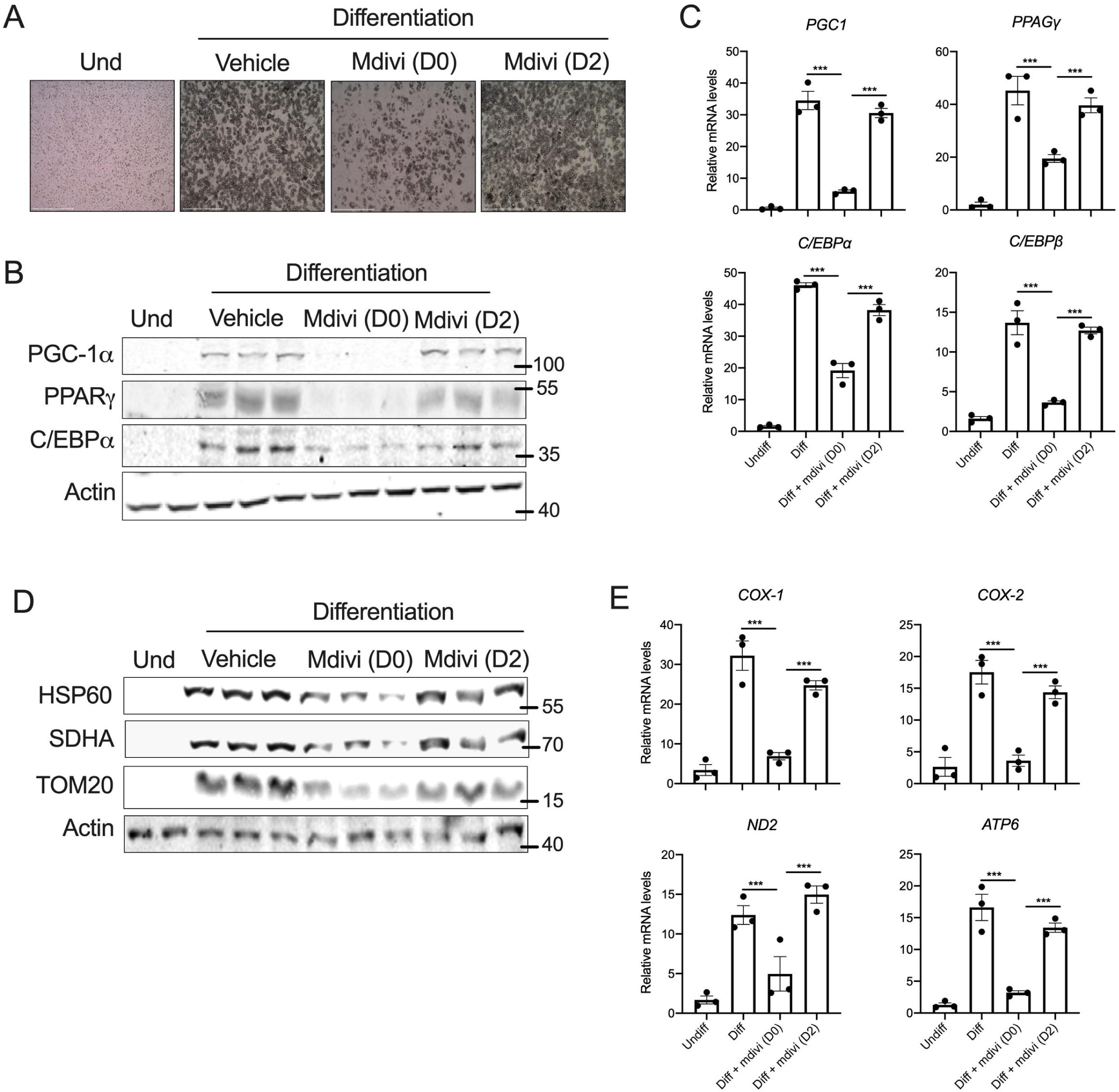
DRP1 is dispensable during the late phase of adipocyte differentiation. Representative images for the SVF cells differentiated to brown adipocytes in the presence or absence of 10μM mdivi-1 added from day 0 (D0) or day 2 (D2) of differentiation and respective treatments continued for additional 6-days. (A) Representative images of the differentiated brown adipocytes. (B) Immunoblots and (C) qPCR analysis for the adipogenesis-related genes. (D) Immunoblots and (E) qPCR analysis for the mitochondria-related genes in the differentiated adipocytes. β-actin was used to normalize the gene expression. The results are presented as SE ± SEM (***p < 0.001). Und-Undifferentiated

## Discussion

Mitochondrial dynamics play a pivotal role in diverse cellular processes, such as adipocyte differentiation, lipid homeostasis, and oxidative metabolism. Adipose tissue mitochondrial dysfunction contributes significantly to the pathogenesis of obesity-associated metabolic diseases (Cedikova et al., 2016; Chouchani & Kajimura, 2019; Dai & Jiang, 2019; Lee et al., 2019). Therefore, it is imperative to identify the mechanisms involved in the biogenesis and functional maintenance of mitochondria in the adipocytes. In the present study, we explored the role of mitochondrial fission protein, DRP1 in the adipose tissue and reveal that DRP1 is highly expressed in BAT and its levels were increased during beigeing and brown adipocyte differentiation. We demonstrate that inhibit of DRP1 during the induction period is sufficient to attenuate adipogenesis and adipogenesis-associated mitochondrial biogenesis. However, DRP1 is dispensable during the post-induction i.e. after induction of differentiation. This is consistent with the previous study, where knockdown of DRP1 in the mature adipocytes did not affect the adipocyte differentiation (Pisani et al., 2018). We demonstrate that DRP1 is essential for the induction of numerous genes involved in the transcriptional programming of adipogenesis. Thus, we propose that DRP1 plays an essential role in adipogenesis.

The adipogenic program is a highly regulated process that involves the formation of mature adipocytes from the precursor cells (Feve, 2005). It is evident from the literature that adipogenesis not only involves the activation of the transcriptional network but also requires mitochondrial remodeling (Wilson-Fritch et al., 2003). During the early phase of adipogenesis, precursor cells are characterized by distinct mitochondrial dynamics and network reorganization, compared to mature adipocytes (Chen & Chan, 2017; Seo, Yoon, & Do, 2018). Our results reveal that the inhibition of DRP1 in adipocyte precursor cells attenuates the early induction of the adipogenic transcriptional factors such as C/EBPα, PPARγ, and PGC-1α.

Adipocyte differentiation is associated with the nuclear translocation of various proteins involved in the regulation of adipogenic transcription factors (Singh et al., 2006). Recent studies have demonstrated a nuclear expression of DRP1 (Chiang et al., 2009), raising the possibility for a transcriptional regulatory role for DRP1 in adipogenesis. However, we were unable to detect the nuclear expression of DRP1 in the precursor or mature adipocytes. DRP1 has been shown to interact with numerous proteins that are involved in cell cycle progression, differentiation, and metabolism (Chou et al., 2012; Ganesan et al., 2019). Therefore, assessing the binding partners of DRP1 will provide novel insights on the mechanism by which cytosolic DRP1 signals nuclear reprogramming of adipocyte transcriptional network. To our knowledge, the role of DRP1 in adipocyte differentiation has never been explored. Our future studies are aimed at understanding the molecular underpinnings of DRP-1 regulation of adipogenesis using mouse models with DRP1 deletion in adipocyte precursor cells.

Obesity and other metabolic diseases are associated with adipose tissue dysfunction and attenuated beigeing and browning efficiencies (Dai & Jiang, 2019). However, the molecular mechanisms are not completely understood. Here, we propose that DRP1 plays an essential role in the commitment of adipocyte precursor cells by regulating the induction of adipogenic transcriptional factors. Further investigation on the expression and post-translational modifications of DRP1 in the adipocyte precursors will provide novel insights on the role of mitochondrial remodeling in obesity and metabolic diseases.

## Experimental procedures

### Animals

C57BL6 mice were purchased from Jackson Laboratories, Bar Harbor, ME. Upon arrival, all the mice were acclimatized in a specific pathogen-free facility in 12-hr day and night cycles and continued on their respective chow diet (D12450J, Research Diets, New Brunswick, NJ). For β3-adrenergic agonist treatment, CL316, 243 was injected intraperitoneally at a dose of 1mg/kg bodyweight once a day for 7-days. All the animal procedures were approved by the University of Pittsburgh Institutional Animal Care and Use Committee.

### Isolation, culture, differentiation and treatment of primary SVF cells

Primary mouse inguinal adipose stromal vascular fractions (SVFs) cells were isolated as described previously (Liu et al., 2017). Briefly, the inguinal white adipose tissue was quickly collected and minced in a petri-dish containing freshly prepared digestion buffer with collagenase type D and dispase II. The tissues were at 37°C for 20-30min with constant agitation. The digestion was neutralized by with DMEM/F12 medium and filtered using 100 uM cell strainer. The filtrate was centrifuged at 500Xg for 5 min at room temperature and the pellet was suspended with the DMEM/F12 medium and further filtered using 40 uM cell strainer and centrifuged at 500Xg for 5 min at room temperature. Finally, the SVF cell pellet was re-suspended and plated in 10CC plate in DMEM/F12 medium supplemented with 10% FBS and 1% penicillin/streptomycin. The preadipocytes were cultured in DMEM/F12 supplemented with 10% FBS and 1% penicillin/streptomycin by maintaining at 37°C. For differentiation, cells were plated in 12-well plate and once cells reach 70-80% confluence (designated as Day0), differentiation was induced with induction media containing 10% FBS, 0.5 mM isobutylmethylxanthine, 125 nM indomethacin, 1 μM dexamethasone, 850 nM insulin, 1 nM T_3_, and 1 μM rosiglitazone (Lu et al., 2018). Two days after induction, cells were switched to maintenance medium containing 10% FBS, 850 nM insulin, 1 nM T_3_ and 1 μM rosiglitazone and continued to maintain for another 4 to 6 days. Maintenance media was refreshed every 48 hours. Details about the Mdivi-1 treatments are detailed below.

### Culture, differentiation and treatment of transformed SVF cells

Unless mentioned as primary SVF cells in the legends, all the experiments using SVF cells were performed using the transformed mouse inguinal SVF preadipocytes cell lines purchased from the Kerafast, Inc. (Boston, MA). Transformed SVF cells were cultured and differentiated as explained above for the primary SVF cells. For DRP1 inhibition, cells were started treated with 10µM Mdivi-1 (Cassidy-Stone et al., 2008) (Cat no: A4472; APExBIO) from day 0 (DO; induction period) or day 2 (D2; post-induction). For Mdivi-1 washout experiment, 10µM Mdivi-1 was added during the induction period (D0) and on D2 the wells were washed and the differentiation process was continued without Mdivi-1.

### Real-Time PCR analysis

Total RNA was extracted using TRIzol reagent (Invitrogen) as per the manufacturer’s instructions. 1μg of total RNA was reverse transcribed using Mu-MLV reverse transcriptase (Promega, Madison, WI). The gene expression was evaluated by Quantitative Real-Time PCR (RT-PCR) using SYBR Green Master mix (Radiant Molecular Tools, Fort Lauderdale, FL) with Quant Studio 3 Station qPCR machine (Applied Biosystems). The results were expressed as fold-change using the 2^-ΔΔCT^ method and β-actin was used as an internal normalization control (List of primer sequences was provided in Table 1).

**Table 1.**
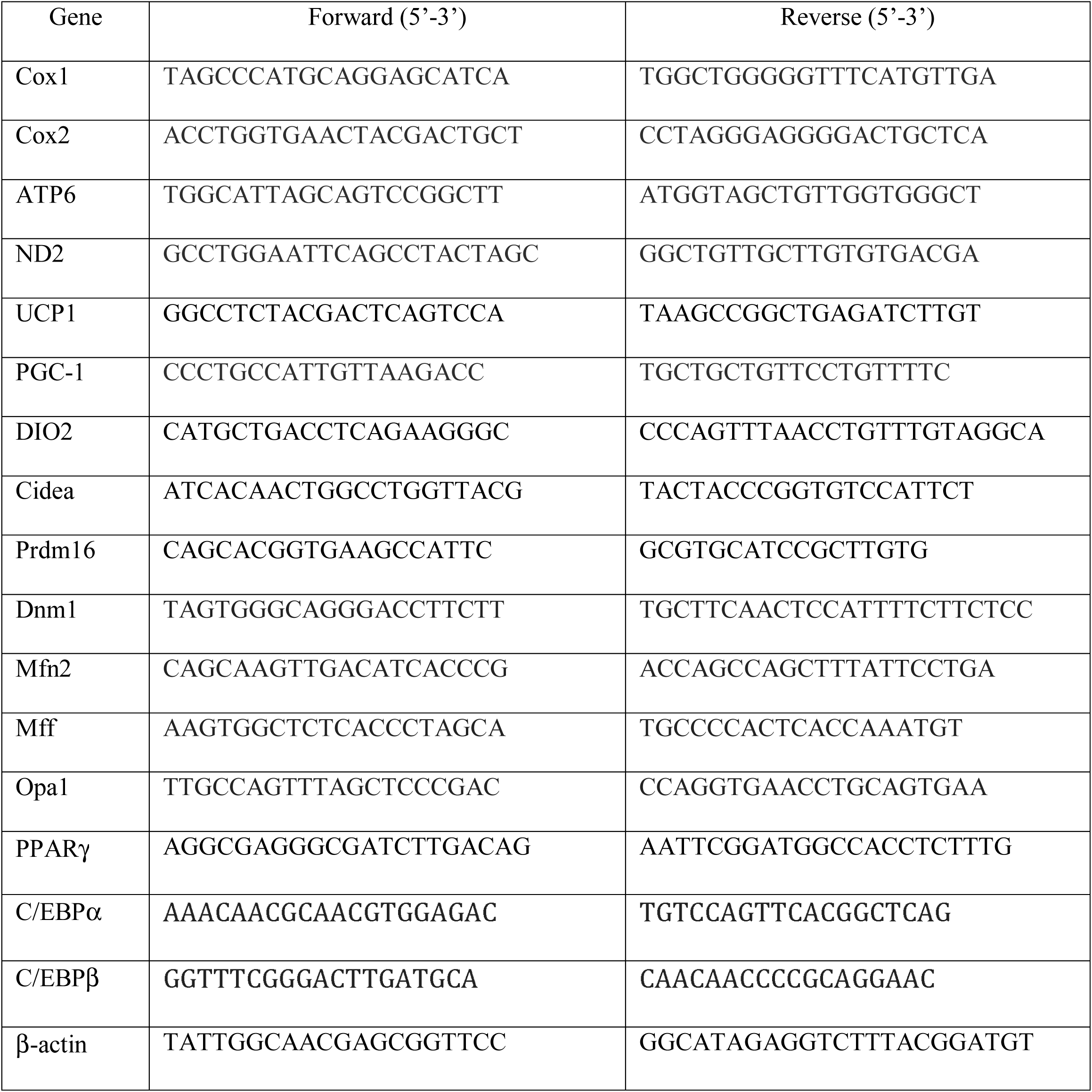
Primers sequence details.

### Western blotting

Protein lysates were prepared using RIPA lysis buffer (50mM Tris pH 7.4, 150mM NaCl, 5mM EDTA, 1% NP-40) containing protease and phosphatase inhibitors (Sigma Aldrich). The protein lysates were separated on 8-12% polyacrylamide gels and transferred to nitrocellulose or PVDF membranes. Membranes were blocked with 5% skim milk in tris-buffered saline containing 0.01% Tween20 and probed with primary antibodies (Table 2) overnight at 4°C. Membranes were then probed with DyLight-conjugated secondary antibodies (Cell Signaling Technology) and visualized using Odyssey CLx Imaging System (LI-COR).

**Table 2.** Antibody catalog number and source.

| **Name** | **Catalogue Number** | **Source** |
| --- | --- | --- |
| Actin | 66009-1-Ig | Proteintech |
| DRP1 | 8570T | Cell Signaling Technology |
| MFF | 84580T | Cell Signaling Technology |
| MFN2 | 11925T | Cell Signaling Technology |
| Opa1 | 80471T | Cell Signaling Technology |
| PGC-1α | ab54481 | Abcam |
| SDHA | 5839 | Cell Signaling Technology |
| Tom20 | 42406S | Cell Signaling Technology |
| UCP1 | PA1-24894 | Invitrogen |

### Nuclear and cytosolic protein lysate preparations

The nuclear and cytosolic protein lysates were prepared by using the NE-PER Nuclear and Cytoplasmic Extraction kit (ThermoFisher Scientific) according to the manufacturer’s instructions.

### Data availability

The data that support the findings of this study are available from the corresponding author upon reasonable request.

### Statistical analysis

The experimental data were analyzed using GraphPad Prism 8 (Version 8.1.2) software. The data are presented as mean ± standard error of the mean (SE). Significance between experimental groups was derived using Unpaired t-test (Two-tailed) for two groups and One-way ANOVA (Tukey’s multiple comparison test) for multiple groups. p-value of <0.05 was considered significant. All the *in vitro* experiments were carried out in triplicate and repeated at least twice.

## Acknowledgements

This work was supported by funding from National Institute of Diabetes and Digestive and Kidney Diseases (DK110537) and Pittsburgh Liver Research Center Pilot and Feasibility grant (P30DK120531) to SKR.

## Conflict of interests

The authors declare no conflict of interest.

## Author contributions

R.G.R. performed experiments, analyzed data and involved in writing the manuscript. D.M, Z.C, and N.B were involved in data acquisition. S.K.R. initiated, designed, and supervised the work, and wrote the manuscript with input from all authors.

